# A Novel Dual And Triple RSVP Paradigm For P300 Speller

**DOI:** 10.1101/441410

**Authors:** Amir Mohammad Mijani, Mohammad Bagher Shamsollahi, Mohsen Sheikh Hassani

## Abstract

**Objective:** A speller system enables disabled people, specifically those with spinal cord injuries, to visually select and spell characters. A problem of primary speller systems is that they are gaze shift dependent. To overcome this problem, a single RSVP paradigm was introduced in which characters are displayed one by one at the center of a screen. In this paper, two new protocols named Dual and Triple RSVP paradigms are introduced and their results are compared against the single paradigm.

**Methods:** In the Dual and Triple paradigms, two and three characters are displayed at the center of the screen simultaneously, therefore holding the advantage of displaying the target character twice and three times respectively, compared to the one-time appearance in the single paradigm. Subsequently, by reducing the number of repetitions in the Dual and Triple paradigms, it is expected that ITR decreases. To compare the results of these three paradigms, three subjects participated in experiments using all three paradigms.

**Results:** The offline results demonstrate an average character detection accuracy of 97% for the single and double protocols, and 80% for the Triple paradigm. In addition, average ITR is calculated to be 5.45, 7.62 and 7.90 bit/min for the single, Dual and Triple paradigms respectively. Results demonstrate an equally good character detection accuracy for the single and Dual paradigms, and a significant increase in ITR in the Dual paradigm compared to the single. The Triple RSVP paradigm demonstrates an almost equal ITR to that of the Dual paradigm, while decreasing character detection accuracy significantly.

**Conclusions:** Results demonstrate that the Dual RSVP paradigm can be recognized as the most suitable approach, by providing the best balance between ITR and character detection accuracy.

**Significance:** This research demonstrates the improved performance of a newly proposed speller system (the Dual RSVP paradigm). By replacing existing methods with this new approach, the performance of speller matrices will be enhanced, and in addition the gaze dependency issue that caused limitations for users suffering from unimpaired oculomotor control will be overcome.

## 1 INTRODUCTION

Brain computer interfaces (BCI) are communication systems that convert the brain’s neural activities into functional commands for external devices (Acqualagna et al., 2013). BCI systems serve a variety of functions such as robot control (Bell et al., 2008), speller systems (Yin et al., 2013) etc. BCI systems are capable of creating direct communication channels to connect the human brain with computers. It can be said that the main purpose of BCI systems is to enable its users, especially those with spinal cord injuries, to communicate with the outside world using only their brain signals. According to this goal description, two main functions are expected from each BCI system; 1) to be able to detect neural activities and understand the purpose of the user and 2) to interpret and translate brain signals into understandable commands for an external device. Amongst the different methods of recording brain signals for use in BCI, Electroencephalography (EEG) is the most commonly used, due to its advantages in providing higher temporal resolution, non-invasiveness, cost efficiency and transportability (Orhan et al., 2012).

Depending on the diverse types of components of EEG signal, BCI systems can be divided into steady state visually evoked potentials (SSVEP)-based (Chen et al., 2017) and event-related potentials (ERP)-based (Zhang et al., 2012) systems, the latter being our area of focus in this paper. Event-related potentials are the brain’s response to an external stimulus or event. ERPs have various components, each of which contains specific information regarding the type of ERP (Sur et al., 2009). One of the most common ERP-based BCI systems is the speller system. Speller systems use an intelligent approach to evoke a specific ERP component, which in most systems is based on the p300 component(Lin et al., 2018). The p300 component has a positive potential that appears with a 300-500 ms delay after the onset of the target stimulus. The p300 component is evoked in an Oddball paradigm when a target stimulus appears among a string of non-target stimuli. This component is usually more evidently visible in central electrodes (C, Fz, Pz) compared to other electrodes (Gonsalvez et al., 2002, Elshout, 2009).

The first ERP-based speller system was introduced by Farwell and Donchin, known as the matrix speller (Farwell et al., 1988). In this protocol, 36 characters, consisting of the letters of the alphabet and single-digit numbers are organized in a 6×6 matrix. In this matrix, the rows and columns are illuminated one by one and randomly, and the user must focus on one of the symbols which is the target-character. The p300 component is expected to be evoked in the subject’s brain signal when the row or column of the target-character is illuminated. By performing numerous repetitions of this experiment and averaging over the target stimuli, the p300 component will be easily observable. The target-character can then be identified by determining the row and column of the evoked p300 component (Farwell et al., 1988).

Although Farwell and Donchin’s protocol was the first paradigm presented for speller systems and was widely used, however, recent studies have shown that target character selection in this paradigm is dependent on eye movement, also referred to as “gaze dependent” (Acqualagna et al., 2010). Therefore, this protocol is not suitable for users suffering from unimpaired oculomotor control. Many studies have been conducted to identify solutions for creating gaze independent speller matrices, which ultimately led to two solutions; 1) changing the type of modality and substituting vision with touch (Höhne et al., 2011) or audio (Brouwer et al., 2010), and 2) changing the movement paradigm. Hex-O-Spell and RSVP paradigms have been suggested in this regard. In the Hex-O-Spell paradigm, characters are divided into six groups, and each group is placed on one edge of a hexagon. This protocol uses a two-step process for target character selection. As the first step, each of the six groups are illuminated one by one and at random, and the subject must focus on the group containing the target character once it is illuminated. This group of characters will be selected as the target group. Then, the characters within the selected target group are placed around a hexagon (one character at each edge) and randomly illuminated, where the target-character can be determined (Treder et al., 2010).

In the RSVP paradigm, characters are randomly displayed one by one at a fast rate on the center of the screen. Target character selection in this method is not gaze dependent (Acqualagna et al., 2010, Orhan et al., 2012). Further changes were applied to the primary RSVP protocol (single RSVP) in order to increase accuracy by performing actions that help better distinguish characters, such substituting upper case letters with lower case and using colored characters (Acqualagna et al., 2011, 2013). Although the RSVP paradigm is considered a new protocol in speller systems and overcomes the gaze dependency issue of the matrix speller, however, displaying only one character at each time slot elongates the experiment duration, which exhausts the user and decreases Information Transfer Rate (ITR). In order to address this time efficiency issue which is present in the single RSVP paradigm, Dual and Triple RSVP paradigms are introduced and utilized in this study. In order to better understand the concept of these two paradigms, another use of the RSVP paradigm (other than speller systems) named “RSVP Image Search” shall be presented. In this paradigm, the goal is to identify the target image from a string of images (Sajda et al., 2003, Huang et al., 2011, Yu et al., 2012, Mohedano et al., 2015). To decrease experiment length and increase ITR in RSVP image search paradigms, Dual and Triple RSVP image search paradigms have been introduced. In the Dual RSVP paradigm presented by Cecotti (Cecotti, 2016), unlike the single paradigm where one image is displayed at a time, two images appear simultaneously at the center of the screen. In this paradigm, the same string of images on the left also appear on the right side, but with a certain delay. For each of the images that are to detected, the user must first gaze at the string of images on the left until the target image appears, and then focus on the string of images on the right until once again observing the target image. To detect the next image target image, the user must once again focus on the left string of images, and this loop continues until all images are observed. A similar approach is followed for Triple RSVP paradigms, as further discussed in this paper (Lin et al., 2017).

**Figure 1.**
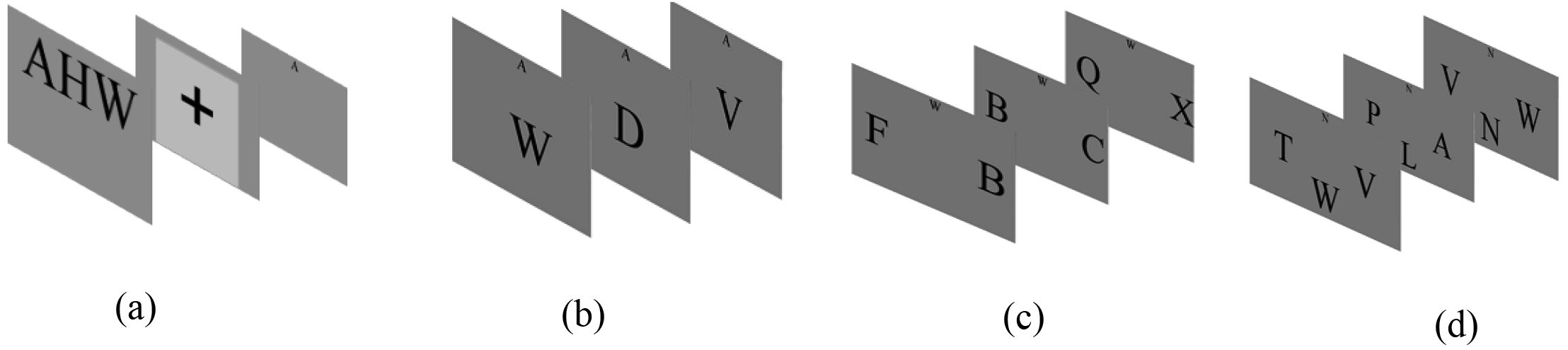
a) Subject preparation, b) Single RSVP paradigm: displaying one character at a time, c) Dual RSVP paradigm: displaying two characters simultaneously in each stimulus, d) Triple RSVP paradigm: displaying three characters simultaneously.

The purpose of this study is to extend the Dual and Triple paradigms presented for image search to speller systems and compare their performance against the single RSVP keyboard paradigm. It is expected that the Dual and Triple RSVP paradigms will improve ITR and reduce experiment duration while maintaining gaze independency.

## 2 METHODS

### 2.1 Participants

Three male subjects within the age range of 25-28 and no previous training in this or any similar experiments participated in this experiment (mean= 25.5, SD= 1.5). Each participant was asked to take part in 3 experiments (one experiment for each paradigm). Subjects passed general medical examinations before participating in the experiments, and showed no symptoms of color blindness, neurological disorders or eye injury. The protocol of these experiments was approved by Iran’s Medical Sciences ethics committee. All participants voluntarily took part in these experiments, and data was recorded in The National Brain Mapping Lab.

### 2.2 Apparatus

EEG signals were recorded using a g.Hlamp (G.Tech company) device. A total of 32 active electrodes were used in our experiments in accordance with the international “10-20” system, and all electrodes were referenced to the right ear. The signal sampling rate was 512 Hz and a digital notch filter was activated at 50 Hz. Stimulation protocols were designed using Psychtoolbox and displayed on a 19.5 inch LED with a resolution of 1366×768 and refresh rate of 60 Hz.

### 2.3 Experiment design and procedure

In this study, single, Dual and Triple RSVP Keyboard paradigms have been designed and experimented on. In all three paradigms, symbols are displayed in black within a grey background. As displayed in figure 1a, to better prepare the subject, the target word appears at the top of the screen for 1 second before the start of each run. Next, the fixation cross appears and vanishes after 1 second, followed by the target character being displayed for 1 second, and finally the stimuli start to appear one after the other (all these steps occur in offline phase).

In the single RSVP paradigm, characters are displayed one at a time at the center of the screen, as demonstrated in figure 1b. In the Dual RSVP paradigm, two characters are displayed at each time frame, as depicted in figure 1c. In other words, two strings are displayed in parallel, where the second string is the delayed version of the first. To create the string and its delayed version, 26 alphabet characters are considered as the primary string. The second string is created by applying a circular shift of 4 units to the first string. An example of two strings created in such a way would look similar to what is demonstrated below. In the Triple RSVP paradigm, three characters are displayed at each time slot, as depicted in figure 1d.

String 1: V P R F J E C S Q T D M O U A Z Y N I H L K G W B X

Shifted string 1: G W B X V P R F J E C S Q T D M O U A Z Y N I H L K

**Figure 2.**
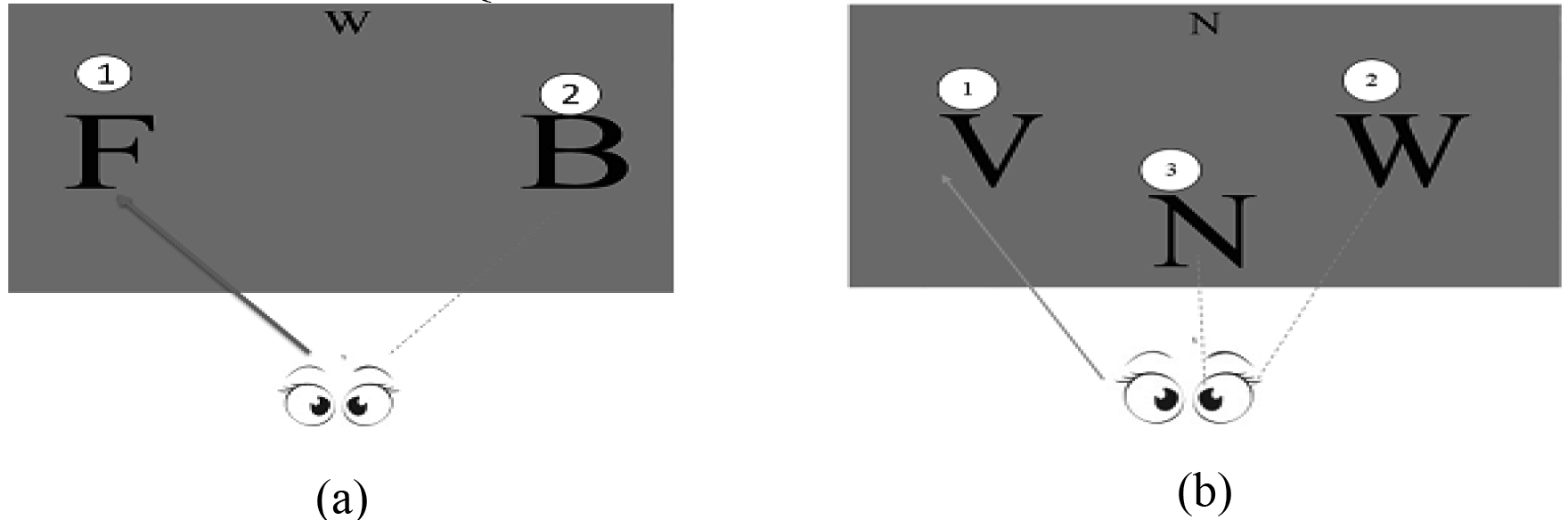
a) Experiment procedure in the Dual RSVP paradigm: the subject is first asked to focus on the left string, and after seeing the target character, shift focus to the right and once again left after character detection, b) Experiment procedure in the Triple RSVP paradigm: the procedure is similar the Dual paradigm, with the addition of third string.

In the Dual RSVP paradigm, the subject is asked to first gaze at the string of characters on the left until the target character appears, then focus on the string of characters on the right. After observing the target character, the user is once again asked to focus on the left string of characters (as demonstrated in figure 2.a). In the Triple RSVP paradigm the goal is to display the target character three times in each repetition, doing so by using three parallel character strings. A similar order applies to the Triple RSVP paradigm, but in this case, after the target character is observed in the right string, the user is asked to gaze at the bottom string. After the appearance of the target character in the bottom string, the user must once again shift attention on the left string. This requires multiple attention shifts between the different strings, creating the possibility of missing the target stimulation while shifting focus from one string to the other in the Dual and triple paradigms (if the strings are parallel with circular shift). For example, if “G” is the target character, after seeing “G” in the left string, the subject will not be able to see the character “G” in the right string. To overcome this problem, the circular shift is replaced with a linear shift, and the three symbols “!”, “.” and “?” are also added to the string. Ultimately, the Dual paradigm contains 29 characters consisting of 26 alphabet letters and three punctuation symbols. A sample string and its delayed version would therefore look similar to this:

1. String 1: UNMRWHYVCKTBZOFLSEAJIPXDGQ!.?
2. Shifted String 1: !.?UNMRWHYVCKTBZOFLSEAJIPXDGQ

To address this issue in the Triple RSVP paradigm, a primary string is created and the second and third strings are created using determined delays. Once again in order to prevent the loss of target character detection, linear delays of 3 and 6 units are applied to the second and third strings, respectively. In addition, digits 1 to 9 are also added to the stimulus characters. Another important factor that must be taken into account in the design of Dual and Triple RSVP paradigms is that no two characters should be repeated together more than once. For example, if two “QX” characters appear together, they must not re-appear together in any of the next iterations. Character strings used in the Dual and Triple RSVP paradigms along with their delayed versions are presented in Tables 1.1 and 2.1. A sample Triple RSVP paradigm character string and its delayed versions can be observed below:

String: UNMRWHYVCKTBZOFLSEAJIPXDGQ123456789
1^st^ shifted string: 456UNMRWHYVCKTBZOFLSEAJIPXDGQ321789
2^nd^ shifted string: 987123UNMRWHYVCKTBZOFLSEAJIPXDGQ645

By using the aforementioned technique to avoid missing target characters, we will have different numbers of characters and different stimulus times for each of the three paradigms. In the single paradigm, 26 characters comprised of letters of the alphabet are used as stimulations. These stimulations are randomly displayed 10 times with stimulus time of 187.5 ms (5.33Hz). Therefore, the time required to spell each character is equal to 187.5×10×26 = 48.75s. In the Dual paradigm, the display of each character is repeated 5 times with a stimulus duration of 250 ms (4Hz). In this paradigm, the delay of the second string is equal to 750 ms. The time required for character detection in this paradigm is 250×29×5 = 36.25s. In the Triple paradigm, each character is displayed 3 times with a stimulation time of 250 ms (4Hz). In this paradigm, the delay of the second and third strings are 750ms and 1500 ms, respectively. The time required to type each character is therefore 250×35×3 = 26.25s. From these calculations, it can be observed that the Dual and Triple paradigms can significantly decrease the time required for character detection.

In the experiments related to each of the protocols in offline mode, subjects are asked to spell 45 characters. Subjects are seated approximately 90 cm from the monitor on a comfortable chair, and are asked to also keep count of the number of times they view the target character. For each protocol, the experiments are performed in a single session and in multiple runs. Each run is comprised of spelling 3 characters, and the subject is given a few minutes of resting time between the runs.

**Table 1.1.**
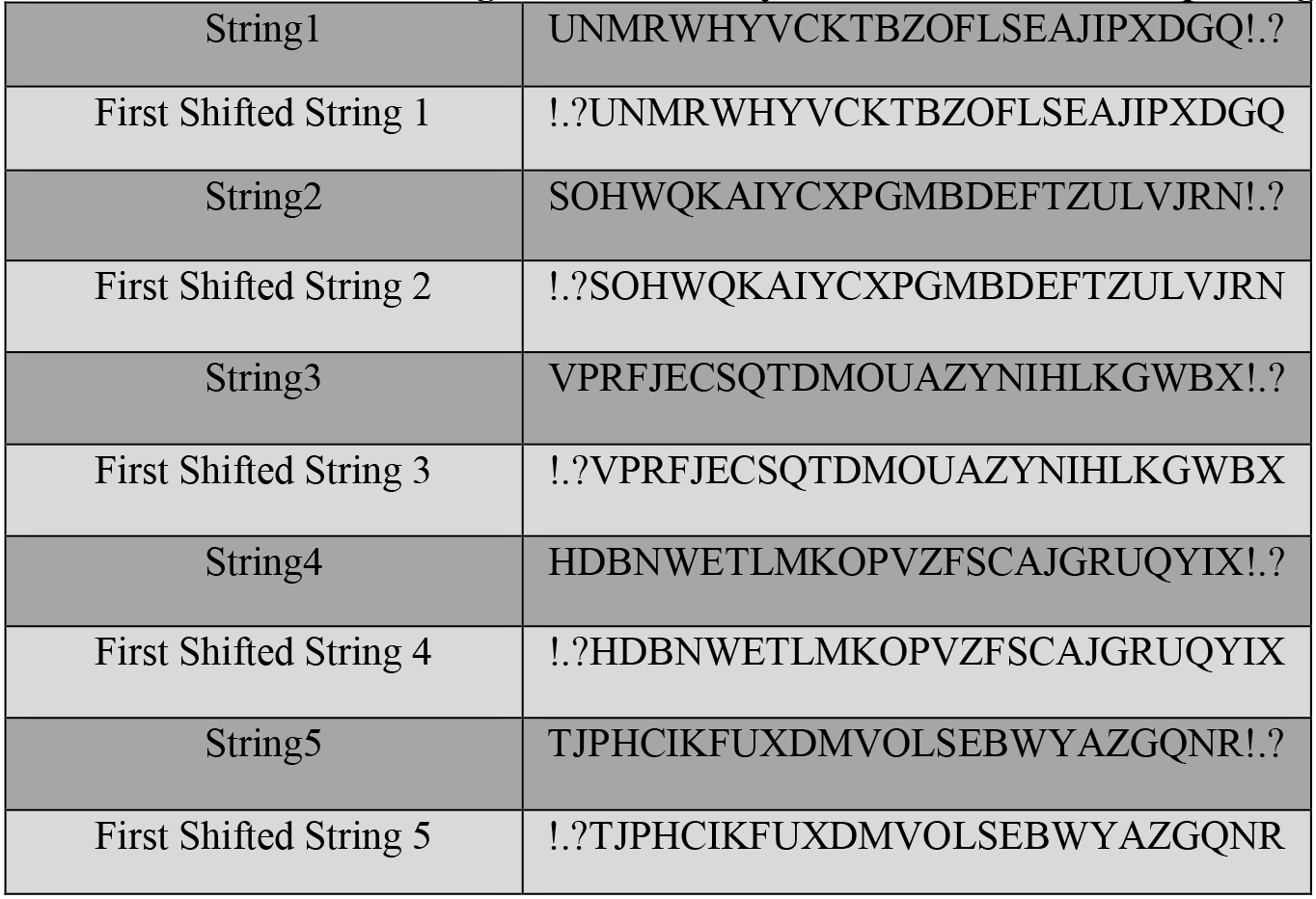
Five character strings and their delayed versions in the Dual paradigm.

**Table 2.1.**
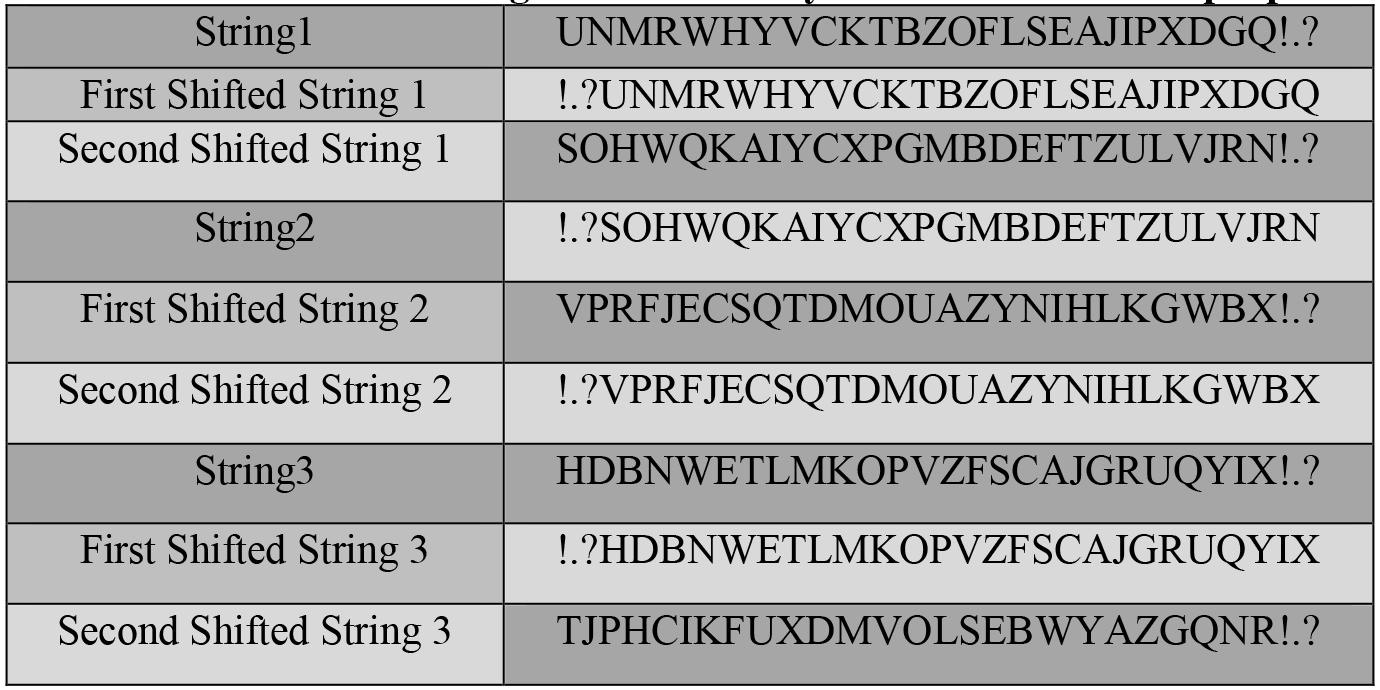
Three character strings and their delayed versions in the Triple paradigm.

### 2.4 Data Analysis

For the purpose of data analysis, all channels were first filtered using a digital band-pass filter in the range of 0.5-40 Hz, and then down-sampled at 128Hz. EEG signals were divided into epochs in the range of 0 to 800 ms after the onset of the stimulation. The mean of 0-100 ms before each stimulation was also deducted from the corresponding epoch for baseline correction. Since the range of recorded EEG signals are approximately between −50 to +50 microvolts, epochs with a max-min range exceeding 100 microvolts were considered artifacts and the entire epoch was removed from the signal in order to remove movement artifacts such as eye movement.

In order to classify target characters from non-target characters, most previous studies have extracted and used temporal features. In addition to temporal samples, wavelet coefficients were also used as features in this study. Amongst the possible features, wavelet transform is one of the most effective for distinguishing between target and non-target stimulations in ERP analysis. To compute the coefficients of wavelet transform, five-octave discrete wavelet transform is applied to each epoch with the mother wavelet “*bior 2.2*” (Tahmasebzadeh et al., 2013). By applying this wavelet transform, the signal is decomposed into coarse detail and coarse approximation. Wavelet coefficients can be obtained by placing approximation coefficients from the last stage and detail coefficients from different stages together (further explained in (Tahmasebzadeh et al., 2013)). Figure 3 demonstrates the average wavelet coefficients for target and non-target stimuli across different trials. As it can be observed, most of p300’s energy in terms of wavelets is amongst a limited number of samples, and these samples demonstrate the most distinction between target and non-target signals. Therefore, only the initial wavelet coefficients (first 50) are used. Temporal samples and retained wavelet coefficients are then placed together to form the feature vector.

Amongst the different available channels, we have used channels F3, Fz, F4, Fc1, Fc2, Cz, Cp1, Cp2, P3, Pz and P4 in our experiments. The selection of these channels was based on two factors, the first being that most similar studies have used the same channels for detecting p300 (Kaper et al., 2004, Rakotomamonjy et al., 2008, Liu et al., 2013). The second reason is that these channels show a stronger p300 on our data compared to the other channels. If we gather the time samples and wavelet coefficients extracted from each epoch using the aforementioned 11 channels, we obtain a 1400-dimension feature vector. Because the dimension of this feature vector is too large, using a dimension reduction method is critical. The Lasso method (Tibshirani, 2011) is applied and the dimension is reduced to approximately 200 to 250 features.

To detect the target character from non-target characters in each experiment, classification is performed using linear RLDA (Acqualagna et al., 2011) and non-linear SVM (Rakotomamonjy et al., 2008) classifiers. Classification is performed separately using each of the named methods and the final results are compared. In our classification method, we use scores instead of class labels to help with target character detection. As previously mentioned, in the Dual RSVP paradigm, two characters are displayed simultaneously in each stimulus. The fact that each stimulus contains two characters at each time frame makes target character identification a complicated issue. Dedicating a score to each of the two characters in every stimulus helps address this issue. Since no two characters will re-appear together, by averaging each character’s score in different appearances, the maximum score determines the target character. It is better to explain the scoring system with an example. Let us say for example that the characters “AB” appear together in a stimulus where “B” is the target character. Theoretically, the score for this stimulus would be 0.9, whereas other stimuli would have a score in the range of 0.1 to 0.2. We pertain this score to both characters “A” and “B”. The challenge now becomes determining which of these two characters are the target character. Since no two characters will re-appear together, we may assume that the next stimuli containing each of these characters would be a different combination of characters, let us assume “BF” and “AX”. In these repetitions, the score of the “BF” stimulus would be approximately 0.9, where as that of “AX” would be 0.2. Therefore, for two repetitions of “B” we obtain scores of 0.8 and 0.9, where as those scores would be 0.2 and 0.9 for “A”. By averaging the scores of each character over different repetitions, the score for “B” would be 0.75, whereas the score for “A” would be 0.3, indicating character “B” is the target stimulus. The same approach is used for target character detection in the Triple RSVP paradigm.

### 2.5 Algorithm Performance Evaluation

The performance of the RSVP-based speller is quantified using ITR and character detection accuracy. In order to accurately estimate classification accuracy, four-fold cross-validation is used. ITR is calculated using the relation below:

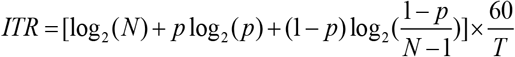

In the above equation, p is character detection accuracy, N is the number of classes and T demonstrates the time required to spell one character (in terms of seconds).

## 3 RESULTS

### 3.1 ERP Analysis

Grand average ERPs for the three protocols on one of the subjects is illustrated in figure 4. In each figure, the blue curve is the ERP response to target stimulus and the red curve is the ERP response to non-target stimulus on the Pz electrode. In figure 4.a, the wave of the p300 component can be observed for the single RSVP paradigm. Figure 4.b demonstrates the ERP waves induced by the Dual protocol. As previously mentioned, target stimulus is displayed twice in each repetition in the Dual paradigm, therefore two p300 components are evoked. The red lines marked in figure 4.b demonstrate the moment of appearance of the target stimulation in the left (0ms) and right (750ms) strings. The ERP wave form produced by the Triple paradigm is also demonstrated in figure 4.c. In this case, three distinct p300 components are evoked. From these wave forms, it can be concluded that the Dual and Triple paradigms are effective in evoking the p300 component.

**Figure 3.**
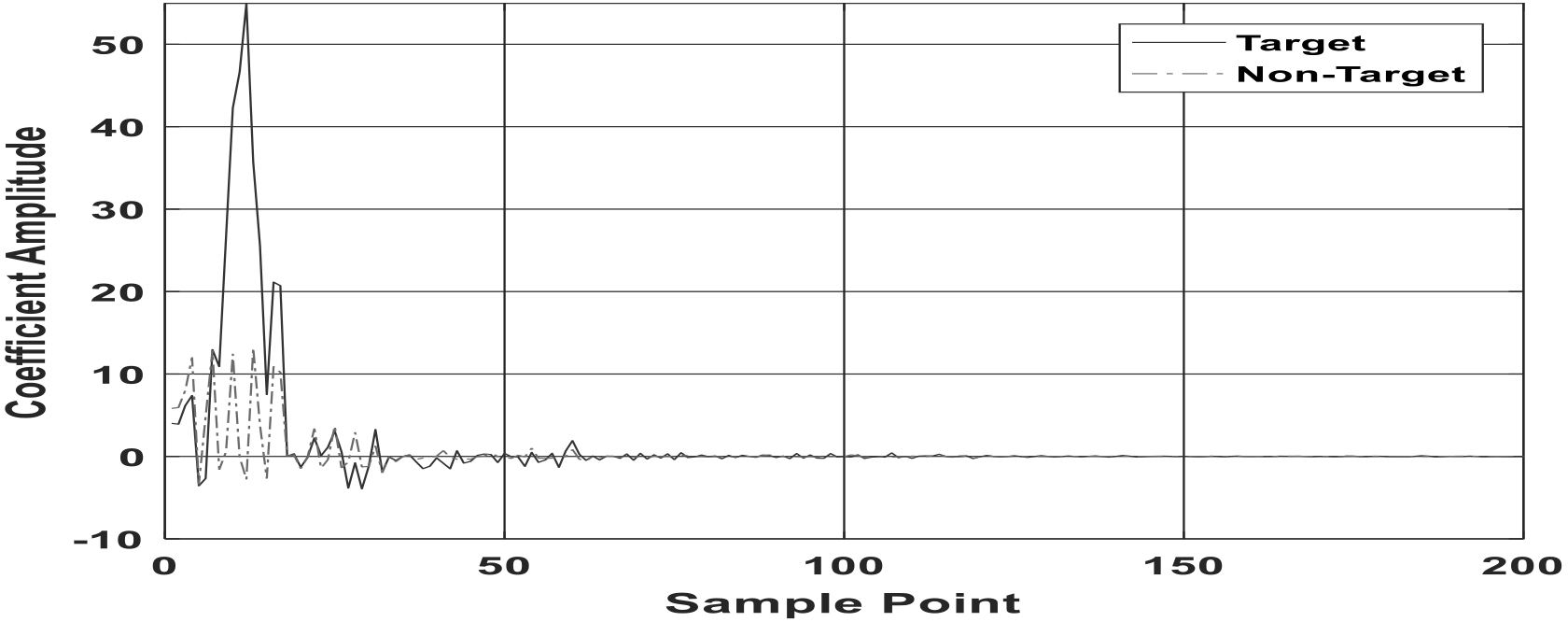
Wavelet transform coefficients obtained using a 5-step discrete wavelet transform. The red curve is the average of target signal coefficients and the blue curve is for the average of non-target signals.

### 3.2 Classification

For classification in offline mode, non-linear ensemble SVM and LDA with covariance shrinkage matrix were used. Character detection accuracy and ITR obtained using these classifiers for the different protocols and all three subjects are reported in table 2. For training the ensemble SVM classifier, three SVM classifiers with Gaussian kernels have been designed, and classifier parameters (c, σ) have been optimized through trial and error. Classification results demonstrate a mean character detection accuracy of 0.97 for the single, 0.97 for the Dual and 0.80 for the Triple paradigms. The ITR for these three protocols is also reported as 5.45, 7.62 and 7.9 bit/min, respectively. The curves for ITR and character detection accuracy for the three subjects have been plotted as a function of different repetitions in figure 5. In these curves, a relatively inverse relationship can be observed between the accuracy rate and ITR. Other than the first iterations, a monotonic decrease in ITR is observed with repetition increase (figure 5b, 5d and 5f), while accuracy increases.

**Table 2.** Character detection accuracy (%) and ITR (bit/min) resulting from the classification of the three protocols.

| Classifier | Ensemble SVM |  |  |  |  |  | RLDA |  |  |  |  |  |
| --- | --- | --- | --- | --- | --- | --- | --- | --- | --- | --- | --- | --- |
|  | Single RSVP |  | Dual RSVP |  | Triple RSVP |  | Single RSVP |  | Dual RSVP |  | Triple RSVP |  |
| Results | ACC | ITR | ACC | ITR | ACC | ITR | ACC | ITR | ACC | ITR | ACC | ITR |
| Sub1 | 0.95 | 5.14 | 0.97 | 7.60 | 0.77 | 7.38 | 0.93 | 4.96 | 0.91 | 6.61 | 0.64 | 5.74 |
| Sub2 | 0.97 | 5.46 | 0.95 | 7.24 | 0.75 | 7.04 | 0.95 | 5.14 | 0.97 | 7.60 | 0.77 | 7.38 |
| Sub3 | 1.00 | 5.76 | 1.00 | 8.04 | 0.88 | 9.28 | 0.97 | 5.37 | 1.00 | 8.04 | 0.88 | 9.28 |
| Mean | 0.97 | 5.45 | 0.97 | 7.62 | 0.80 | 7.90 | 0.95 | 5.15 | 0.96 | 7.41 | 0.76 | 7.46 |
| S.D | 0.02 | 0.31 | 0.02 | 0.40 | 0.07 | 1.20 | 0.02 | 0.20 | 0.05 | 0.73 | 0.12 | 1.77 |

## 4 DISCUSSION

After obtaining the results from our experiments, we aim to compare the three protocols in terms of character detection accuracy and ITR. Average ITR for the single, Dual and Triple RSVP paradigms is observed to be 5.45, 7.62, and 7.90, with a mean character detection accuracy of 0.97, 0.97 and 0.80, respectively. As it can be observed, although the average character detection accuracy is higher in the single protocol compared to the triple, however, average ITR in the Triple protocol is much higher.

As mentioned, in the single paradigm the target character appears once in each repetition, so with 10 repetitions, the target character appears 10 tines. In the Dual paradigm the target character appears twice in each repetition, and therefore it appears 10 times in 5 repetitions. In the Triple paradigm, the target character appears 9 times in 3 repetitions, as it appears 3 times in each repetition. Since in the single, Dual and Triple paradigms, 10, 10 and 9 p300 components are evoked respectively, it was expected that the Dual and Triple paradigms show a much better ITR while maintaining accuracy. However, the results demonstrate that the high character detection accuracy of the single paradigm is sustained in the Dual paradigm but has decreased in the Triple paradigm. The increasing trend in ITR has however fulfilled the expectations, as ITR in the Triple paradigm increases significantly compared to the single paradigm. Yet, the significant advantage of the Triple paradigm is that the time it requires for 9 p300 component stimulations is almost half of the single paradigm (for 10 p300 component stimulations).

Another conclusion that can be made by comparing the results of the three protocols is that in the Dual paradigm, in addition to maintaining accuracy, ITR increases significantly compared to the single paradigm. The Triple paradigm however results in a significant accuracy drop compared to the Dual paradigm, while not increasing ITR by a significant amount. Therefore, by considering these results, the Dual paradigm can be introduced as the most successful.

In BCI systems and especially speller systems, a trade-off exists between character detection accuracy and ITR. As character detection accuracy increases with more repetitions, ITR decreases, and vice versa (as results from the Single Dual and Triple paradigms indicate). Therefore, depending on which of these parameters is of more importance for the purpose of a specific experiment, the other must be sacrificed for the improvement of that parameter.

An issue with the Dual and Triple paradigms is that after being displayed in the left string, the target character is re-displayed in the right (and also bottom for the Triple paradigm) string after 3 (and 6 for the bottom string) characters. Since the users are asked to shift focus on the next strings after viewing the target character in the left string, and the delay between the two strings is only 3 characters, the subject will see the target character as soon as shifting focus to the next string. After a few repetitions, this trend in target character appearance in the other two strings will become discoverable. This predictability may affect the Oddball nature of the experiment. Considering the results reported in table 2, we may claim that the observed decrease in the p300’s amplitude in the Dual and Triple paradigms does not disrupt the accuracy of the system and these paradigms are still effective. As depicted in the ERP curves for the Dual (figure 4.b) and Triple(figure4.c) paradigms, the amplitude of the p300 component in the second and third strings must be smaller than the amplitude of the p300 component in the first string.

We may also extend the findings from these experiments to a quadruple RSVP paradigm. In this protocol, 4 characters would be displayed at the center of the screen simultaneously. Although this protocol will decrease the experiment time, however, the fourth p300 component amplitude will most likely undergo a significant decrease, and despite adding complexity to this protocol, it will probably not increase the system’s accuracy.

**Figure 4.**
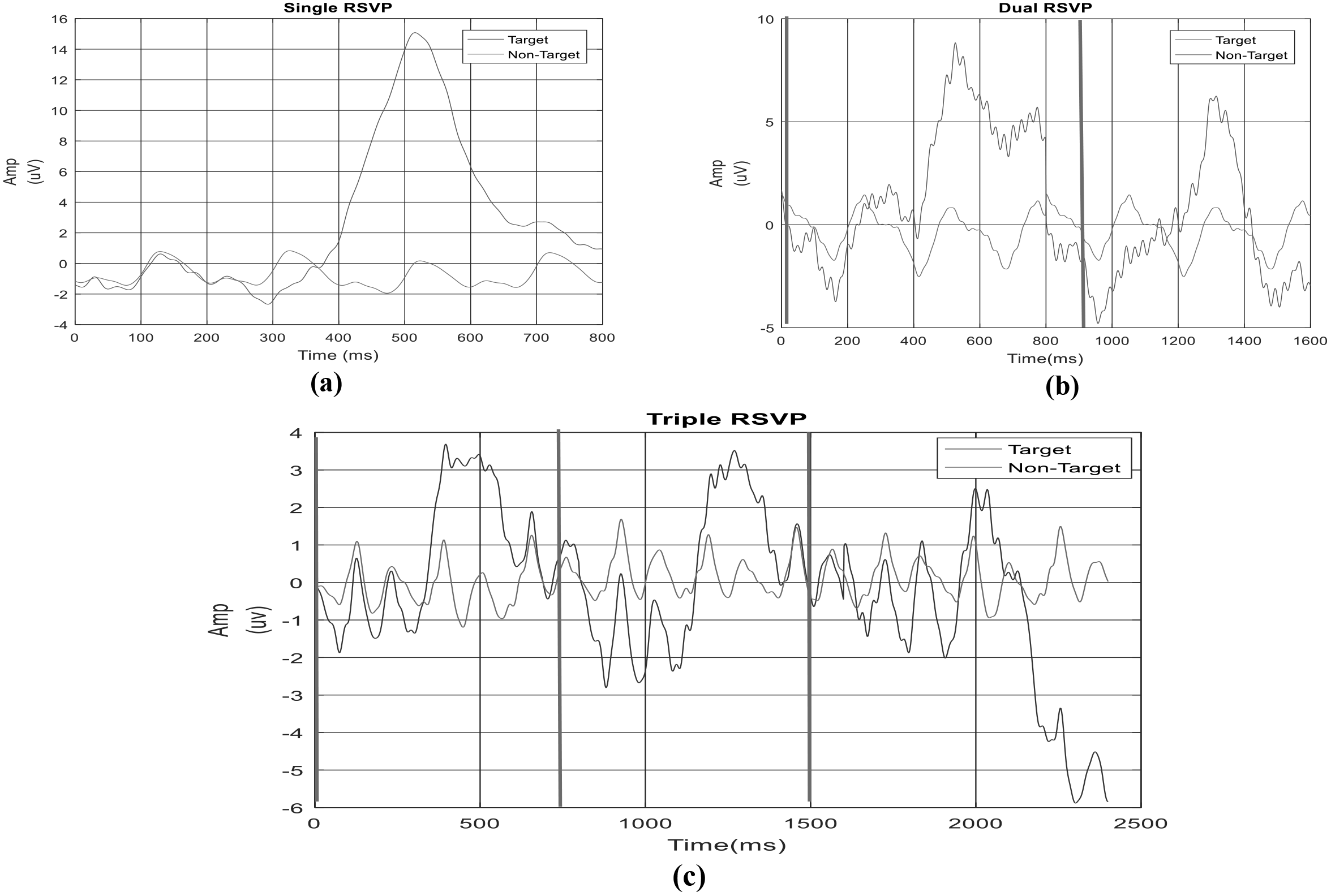
Target and non-target ERP component curves for the three protocols; a) single b) Dual and c) Triple RSVP. In each plot, the blue and red curves correspond to target and non-target p300 component detection, respectively.

Overcoming the gaze dependency problem in the previous speller matrices was the main motivation which led to the development of the primary RSVP paradigm. In this study, we have extended this approach and presented Dual and Triple paradigms to improve character detection accuracy and ITR while maintaining gaze independency. Although in the Dual and Triple paradigms the user is asked to change focus between 2 or 3 different characters, but since the characters are placed close to each other and at the center of the screen, target character detection in these paradigms is not gaze dependent. Through gathering feedback from the users after each experiment, it was found that the Dual and Triple paradigms hold an advantage in terms of user comfort in focusing on the stimuli and character detection compared to the single paradigm. Since the rate of stimulus display in the single paradigm is much faster compared to the Dual and Triple paradigms, all users believed looking at the stimuli in the single paradigm required higher focus and thus their eyes felt more exhausted.

Lastly, the major limitation of this study is regarding our test subjects, where we were only able to recruit 3 male subjects. This number of participants may be too small, and also too specific, considering they were all young male students. As future work, we suggest increasing the number of participants in a more various population.

## 5 CONCLUSION

In this study, by comparing the results of the three single, Dual and Triple RSVP paradigms, we aimed to determine the best approach in terms of ITR and character detection accuracy. By conducting some experiments, we obtained an average character detection accuracy of 97% for the single and double protocols, and 80% for the triple paradigm. In addition, average ITR was calculated to be 5.45, 7.62 and 7.90 bit/min for the single, Dual and Triple paradigms respectively. Results demonstrated an equally good character detection accuracy for the single and Dual paradigms, and a significant increase in ITR in the Dual paradigm compared to the single. The Triple RSVP paradigm demonstrates an almost equal ITR to that of the Dual paradigm, while decreasing character detection accuracy significantly. Therefore, the Dual paradigm can be introduced as the most suitable approach.

**Figure 5.**
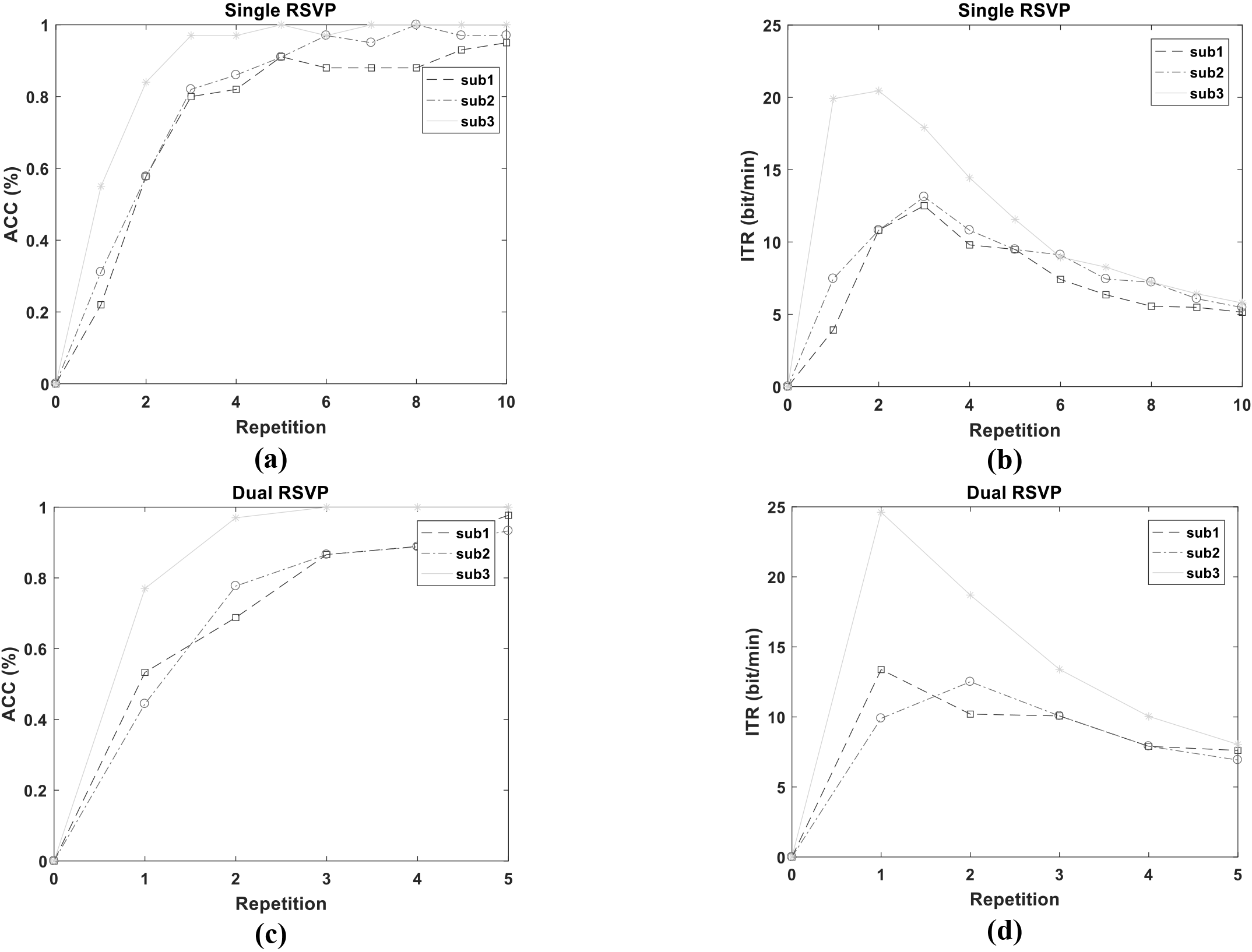
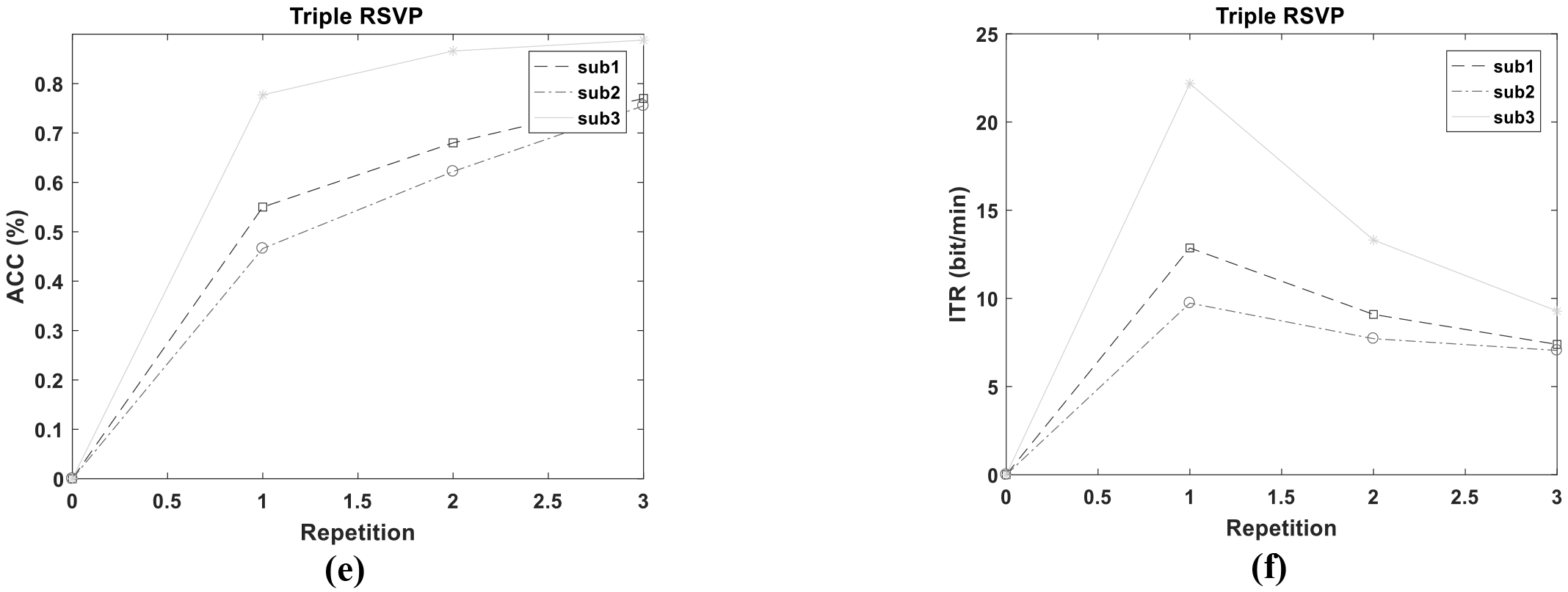
Character detection accuracy and ITR as a function of number of repetitions in the single (a, b), Dual (c, d) and Triple (e, f) paradigms.

## Acknowledgements

We would like to thank The National Brain Mapping Lab for helping us in our data recording process.

## Conflict of Interest(s)

None declared.

